# NAD metabolism mediates a sex-specific protection against early life infections

**DOI:** 10.64898/2026.10.19.760092

**Authors:** Björn Klabunde, Karoline Aasmul Olsen, Malene Spiegelhauer, Emilie Dalbram Kellezi, Adela Memushaj, Anna Hammerich Thysen, Nguyen Phuoc Long, Susanne Brix, Casper Pedersen, Klaus Bønnelykke, Bo Chawes, Jonas Treebak, Ole Bæk, Duc Ninh Nguyen

## Abstract

Sex influences early-life immunity and infection outcomes, yet how metabolic mechanisms regulate these effects remain unclear. As nicotinamide adenine dinucleotide (NAD) can support mitochondrial metabolism and immune function, here we explore how NAD metabolism contributes to sex-specific protection against infections in early life. First, in two independent cohorts of 700 and 386 children, we found that females at birth had higher blood NAD levels than males, and strikingly higher birth NAD levels were associated with reduced infection burdens during the first 3 years of life in females, but with tendency of increased burdens in males. At 18 months, plasma nicotinamide associated with sex-specific cytokine responses to antigen challenges and showed a stronger positive association with tricarboxylic acid cycle (TCA) activity in females. Next, we verified the causal effects of NAD metabolism on infection protection by testing the supplementation of NAD precursor nicotinamide riboside (NR) in infected newborn piglets, used as a translational model of infection-sensitive infants. NR treatment improved survival, bacterial clearance, and disease tolerance only in females, but not in males. Mechanistically, NR increased levels of hepatic TCA cycle metabolites fumarate and malate, and enhanced hepatic inflammatory, hormonal, and mitochondrial-associated transcriptional programs, again only in females. Finally, the sex-specific immune functions mediated by NR were also validated in blood and immune cells *in vitro*. Collectively, we identified NAD metabolism as a sex-dependent immunometabolic mediator in early-life infection. Our findings suggest the potential of sex-stratified nutritional interventions in infection-sensitive infants.

## Introduction

Early life and childhood are periods of increased susceptibility to infections, with a median of 16 infectious episodes up until the age of three.^1,2^ Most common childhood infections are self-limiting and resolved without apparent long-term consequences but do cause substantial health care costs. However, a higher burden of infections in early life is associated with diminished lung function and higher antibiotic use later in childhood.^3^ In vulnerable newborns, especially in those born preterm or growth restricted, infections may cause uncontrolled pro-inflammatory signaling and substantial tissue damage, which may ultimately lead to death.^4–8^ As such prevention of early life infections, or interventions to dampen infection severity, are critical to ensure long term health in children.

It is known that sex affects infection risk and infection outcomes, also in children. In infancy and childhood, females experience higher incidence and severity of several viral and intracellular bacterial infections, whereas male sex is a consistent risk factor for bacterial infections and neonatal sepsis.^9–11^ Mechanistically, females tend to mount stronger immune responses to similar stimuli than males. This has been explained by higher expression of X-linked immune genes and early-life differences in sex hormones, which result in higher expression of genes related to toll-like receptor (TLR) signaling pathways in females.^12,13^ This improved immune activation may lead to more symptomatic infectious episodes, but eliminate pathogens before they progress to severe disease and can come at the cost of greater immunopathology and higher risk for autoimmune disorders later in life.^12–14^

Immune responses to infections require rewiring of metabolic pathways, not only in immune cells but also in tissues exposed to pathogens or inflammatory mediators. Mitochondrial metabolism is central to the resolution of inflammation and maintenance of normal tissue function during infections. Dysregulation of mitochondrial metabolism is associated with sepsis progression, a state of uncontrolled pro-inflammatory activity, both in children and adults. Amongst the metabolic pathways dysregulated, are those related to nicotinamide adenine dinucleotide (NAD) metabolism.^14–16^ NAD acts as the main electron transporter in mitochondrial energy metabolism and a key cofactor for a wide range of enzymatic reactions. We have previously shown that reduction in NAD metabolism is associated with decreased mitochondrial oxidative phosphorylation and poorer infection outcomes in experimental early life bacterial infections.^17,18^ We have likewise shown that increased activity of the mitochondrial tricarboxylic acid (TCA) cycle is connected to both reduced risks of common childhood infections and infection severity in animal models.^19^ Increased levels of NAD were previously linked to improved infection outcomes.^14,20^ NAD is generated *de novo* from tryptophan and during use can be salvaged from Nicotinamide (NAM) or the related molecule Nicotinamide Riboside (NR), a form of vitamin B3. Supplementation with NR is a well proven way to increase intracellular NAD levels.

With this background, we hypothesized that sex differences in immune responses and infection risk or severity are mediated via NAD metabolism. We further hypothesized that boosting NAD levels during severe infection could alter disease outcomes in a sex-dependent manner. First in a human birth cohort, we found significantly higher blood NAD levels at birth in females compared to males. Further, there was a significant interaction between sex and infection risk mediated by NAD, with NAD levels associated with lower infection risk in females but higher risk in males. These results were verified in an independent birth cohort. Later in life, plasma levels of NAM correlated with improved immune responses and TCA cycle metabolism only in females. Then we explored the impact of boosting NAD during an experimental early life infection. Supplementation of NR to infected newborn piglets enhanced immune response, bacterial clearance and infection survival only females but not males, while increasing hepatic NAD levels more in males. Likewise, NR supplementation increased hepatic levels of TCA cycle metabolites and upregulated genes related to OxPhos only in female animals. These consistent findings across clinically relevant settings provide proof-of-concept for how NAD metabolism modulates immunometabolic response to infection in sex-dependent manner and suggests that supplementation of NR to female children may prevent or treat infections.

## Material and Methods

### Human cohort data

#### Study population

We used data from the Danish population-based COPSAC_2010_ mother–child cohort, which has been described in detail previously, including recruitment procedures and baseline characteristics (Chawes et al., 2016). The cohort was embedded in two randomized controlled trials. Pregnant women were randomized in a 2 × 2 factorial design to receive either fish oil capsules containing 2.4 g/day n-3 long-chain polyunsaturated fatty acids or placebo from pregnancy week 24 (ClinicalTrials.gov: NCT00798226). A subgroup of women was additionally randomized to high-dose vitamin D supplementation of 2400 IU/day or standard-dose supplementation of 400 IU/day (ClinicalTrials.gov: NCT00856947). The study was approved by the Danish Ethics Committee (H-B-2008-093) and the Danish Data Protection Agency (2008-41-2599). Written informed consent was obtained from all participants, and the study was conducted in accordance with the Declaration of Helsinki and the Belmont Report.

During the first 3 years of life, infections were prospectively captured through daily diaries and acute visits to the COPSAC clinic. Recorded infection outcomes included pneumonia, tonsillitis, acute otitis media, gastroenteritis, cold symptoms, and fever episodes. Diagnoses were verified by COPSAC physicians at scheduled and acute visits using parental interviews, clinical records, and registry information.

#### NAD and NAM measurements

NAD levels were quantified in relative abundance in neonatal dried blood spot samples collected at birth using nuclear magnetic resonance (NMR)-based metabolomics. At 6 and 18 months of age levels of NAM in plasma were quantified by LC-MS performed by Metabolon, Inc. (Durham, NC, USA).^21,22^ Both metabolites were identified at level 1 according to the established MSI reporting standards.^23^

#### Immune profiling at 6 and 18 months

At 6 months of age, plasma levels of inflammatory markers were measured. At 18 months of age, heparinized whole blood was collected and processed within 4 h for immune cell phenotyping and ex vivo immune stimulation, likewise described in detail elsewhere.^24^ Immune cell subsets were quantified by flow cytometry, including granulocytes, neutrophils, eosinophils, B cells, T cells, CD4 and CD8 T cells, regulatory T cells, γδ T cells, invariant natural killer T cells, monocyte subsets, dendritic cell subsets, and natural killer cell subsets. Functional immune responses were assessed by stimulating whole blood for 24 h at 37°C and 5% CO with lipopolysaccharide (LPS), peptidoglycan (PGN), Poly(I:C), R848, HDMAPP, or staphylococcal enterotoxin B (SEB). Cytokine and chemokine concentrations were measured in stimulation supernatants using MesoScale Discovery multiplex assays. Values below the detection limit were assigned half the lower limit of detection. Cytokine stimulation ratios were calculated by dividing stimulated concentrations by unstimulated concentrations, followed by z-score transformation prior to analysis.

### Animal and *in vitro* studies

#### Cultivation of S. epidermidis

*S. epidermidis* (WT 1457, isolated from a septic infant) was cultured in brain heart bouillon overnight at 37°C / 180 rpm. On the day of the infection, a day culture was inoculated from the overnight culture and incubated until the early logarithmic phase (OD600 = 1). Bacterial density was determined using a spectrophotometer. Bacteria were centrifuged and suspended in sterile saline to a final OD_600_ = 1, equivalent to 3 × 10^8^ CFU/ml. Piglets were inoculated with 10^9^ CFU/kg body weight by arterial infusion over 3 min. Control piglets were infused with a corresponding volume of sterile saline.

#### Animal experimental procedures

All animal studies and experimental procedures were approved by the Danish Animal Experiments Inspectorate under license number 2020-15-0201-00520. These approvals are in accordance with the EU Directive 2010/63, which governs the legislation for the use of animals in research. Details in animal procedures were performed as described before^31,32^. Briefly, crossbred preterm piglets (Landrace × Large White × Duroc) were delivered by cesarean section at day 106 of gestation (∼90% of term) from healthy sows. Immediately after birth, the piglets were transferred to a controlled intensive care unit, where they were housed individually in ventilated incubators preheated to 37°C with an initial oxygen supply of 1–2 L/min. Piglets were resuscitated using Doxapram and Flumazenil (0.1 mL/kg, intramuscular injection), tactile stimulation, and positive airway pressure ventilation until respiratory stability was achieved. Each piglet was equipped with a vascular catheter inserted into the dorsal aorta via the transected umbilical cord, allowing for parenteral nutrition (PN) administration, bacterial inoculation, and blood sampling. Approximately 2 h postpartum, all piglets were randomized into experimental groups based on birth weight and sex.

#### Experimental groups and infections

Piglets received intraarterial PN formulations and were challenged with live *S. epidermidis* (SE) via intraarterial infusion over 3 min. In a first experiment, NR-treated piglets received an initial bolus of 200 mg/kg NR in 2 ml saline (N=14; 8 females, 6 males). An additional 200 mg/kg was administered via PN. Controls received an equal volume of saline (N=18; 11 females, 7 males). In a second experiment, NR treated piglets received 50 mg/kg NR as a bolus and an additional 50 mg/kg over PN (N=24; 11 females, 13 males), with controls receiving saline (N=23; 12 males, 11 females).

#### Clinical monitoring and humane endpoints

Throughout the experiment, piglets were continuously observed for clinical signs of sepsis. Animals reaching predefined humane endpoints, including an arterial blood pH of ≤7.1, deep lethargy, discoloration, and tachypnea, were euthanized for tissue collection and categorized as non-survivors.

#### Blood sampling and analysis

Blood samples were collected at 3, 6, 12 h post-inoculation and at euthanasia via the arterial catheter for blood gas analysis, hematology, and plasma storage for cytokine measurements. At euthanasia, serum and plasma samples were collected for biochemical and metabolic analyses. To determine bacterial counts in blood, jugular vein punctures were performed at 3-and 6-h post-infection and at euthanasia. Blood gas analysis was performed using the GEM Premier 3000 (Instrumentation Laboratory, USA). Hematological assessments were performed using the ADVIA 2120i Hematology System (Siemens, Germany). For cytokine analysis, plasma concentrations of TNF-α, IL-6, and IL-10 were quantified using porcine-specific DuoSet enzyme-linked immunosorbent assays (ELISA) (R&D Systems, USA) according to manufacturer’s instructions. Serum biochemistry evaluations, including liver and kidney function markers, were conducted using the ADVIA 1800 Chemistry System (Siemens, Germany).

#### Euthanasia and tissue collection

At humane endpoints or scheduled euthanasia times, all piglets were deeply anesthetized with a Zoletil mixture (0.1 ml/kg), which included Zoletil 50 (125 mg tiletamine, 125 mg zolazepam), xylazine (6.25 ml xylazine 20 mg/ml), ketamine (1.25 ml ketamine 100 mg/ml), and butorphanol (2.5 ml butorphanol 10 mg/ml) and were subsequently euthanized with an intracardiac injection of barbiturate. The liver was collected during necropsy and snap-frozen in liquid nitrogen, followed by storage at -80°C until liver transcriptomic analysis.

#### RNA extraction and transcriptomic analysis

Liver tissues collected at euthanasia were used for transcriptomic analysis. RNA was extracted using the RNeasy Mini Kit (QIAGEN, USA), and whole-transcriptome shotgun sequencing was performed for gene expression profiling. Library preparation and sequencing were conducted by NOVOGENE (Cambridge, UK). RNA-Seq libraries were generated from 1000 ng of total RNA using the VAHTS mRNA-Seq V3 Library Prep Kit for Illumina (Vazyme, Nanjing, PRC) and sequenced on the Illumina NovaSeq 6000 platform, producing 150 bp paired-end reads. Raw reads underwent quality control and adapter trimming using TrimGalore (Babraham Bioinformatics, Cambridge, UK). The cleaned reads, averaging approximately 26 million per sample, were aligned to the porcine reference genome (Sscrofa11.1) using Tophat2. Gene annotation was obtained from Ensembl (release 99), and a gene-count matrix was generated using htseq-count.

For differential gene expression (DEG) analysis, weakly expressed genes (less than 10 copies in a maximum of 3 samples) were filtered out, and only protein-coding genes were included. DESeq2 package (version 1.38.3)^25,26^ was used to identify DEGs across the following comparisons: NR versus control (all animals), NR vs control (females only), NR vs control (males only), NR female vs NR male, control female vs control. Results were adjusted for litter. Fold-change values were estimated using the lfcShrink function with ashr, and an adjusted false discovery rate (FDR) of 0.05 was applied to determine statistically significantly enriched transcripts. Volcano plots depicting total gene expression were visualized using the EnhancedVolcano package (version 1.14.0)^27^.

Gene set enrichment analysis (GSEA) was performed to identify significantly altered biological pathways using the clusterProfiler package (version 4.0.2)^28^. Genes were ranked by log2 fold-change values before pathway analysis. Pathway annotations were obtained from the Kyoto Encyclopedia of Genes and Genomes (KEGG) for *Sus scrofa*. An FDR threshold of 0.05 was used to define significantly enriched pathways. The normalized enrichment score (NES) was determined, with positive and negative values indicating upregulated and downregulated pathways, respectively. Heatmaps depicting the relative expression of genes within enriched pathways were generated on Z-score adjusted samples, using the ComplexHeatmap package (version 2.15.1)^29^.

#### Targeted Metabolomics

Targeted LC-MS analysis was performed on liver extracts prepared in 80% methanol containing an internal standard mix of citric acid-¹³C, malic acid-¹³C, fumaric acid-d, succinic acid-d, α-ketoglutaric acid-d, D-3-hydroxybutyrate-¹³C, and 2-deoxy-D-glucose-6-phosphate. Pulverized liver tissue was weighed into cryotubes, mixed with extraction solvent at 200 µL per 10 mg tissue, and homogenized with three 2.8 mm ceramic beads using a TissueLyser II at 30 Hz for three 30 s intervals. Samples were then sonicated in an ice bath for 5 min, incubated on ice for 30 min to precipitate proteins, and centrifuged at 11,292 × g for 3 min at 4 °C. Supernatants were transferred to LC vials, with aliquots reserved for 3-nitrophenylhydrazine derivatization and sugar phosphate derivatization, and stored at −80 °C until further processing. Extraction blanks and pooled quality control samples were prepared in parallel.

TCA cycle metabolites were analyzed following 3-nitrophenylhydrazine derivatization to improve detection of carboxylic acid-, ketone-, and aldehyde-containing metabolites. Fresh derivatization reagents were prepared as 200 mM 3-nitrophenylhydrazine in 50% methanol, 120 mM EDC in 50% methanol containing 6% pyridine, and 0.05 mg/mL BHT in 30% methanol. For each sample, 20 µL of 3-nitrophenylhydrazine solution and 20 µL of EDC solution were added sequentially, vortexed, and incubated for 1 h at room temperature. The reaction was quenched with 40 µL BHT solution, after which samples were vortexed and stored at −20 °C until analysis. Blanks, pooled samples, and calibration standards containing the target compounds were derivatized using the same procedure.

LC-ToF MS analysis was performed using an Agilent 1290 Bio UPLC coupled to a Bruker Impact time-of-flight mass spectrometer. Samples were randomized before analysis and injected at 2 µL onto a Waters HSS T3 column, 2.1 × 100 mm, 1.8 µm, maintained at 30 °C. Metabolites were separated at 0.35 mL/min using mobile phases consisting of 0.1% formic acid in water, solvent A, and 0.1% formic acid in acetonitrile, solvent B. The gradient was 5% to 100% B from 0–12 min, held at 100% B from 12–12.5 min, returned to 5% B from 12.5–13 min, followed by a 1.5 min equilibration. The mass spectrometer was operated in negative ion mode with a VIP-HESI source at 3000 V, a scan range of m/z 50–1000, and a spectral acquisition rate of 2 Hz. The nebulizer pressure was 2.0 bar, dry gas and probe gas flow rates were 8 and 4 L/min, respectively, and temperatures were 220 °C and 400 °C. Sodium format cluster ions acquired during the first 0.3 min of each run were used for calibration. Methanol solvent blanks were injected after every six samples to monitor carryover, and samples were maintained at 4 °C during analysis. Data were processed in Skyline v.24.1.0.199 using molecule transition lists containing compound names, chemical formulae, adducts, and explicit retention times. Target retention times are listed in **Supplemental Table 1**.

#### NAD cycling assay

Tissue NAD and NADH content was measured using an alcohol dehydrogenase–coupled enzymatic cycling assay.^30^ Frozen tissue was homogenized in 400 µL of 0.6 M perchloric acid and centrifuged to remove insoluble material. The supernatant was diluted in 100 mM sodium phosphate buffer, pH 8.0, and mixed in equal volume with a freshly prepared cycling reagent containing ethanol, alcohol dehydrogenase, diaphorase, nicotinamide, flavin mononucleotide, and resazurin. The increase in resorufin fluorescence was monitored kinetically at 540–544 nm excitation and 580 nm emission. NAD concentrations were determined from the linear reaction rates relative to an NAD standard curve and normalized to tissue mass or protein content.

#### Cell culture experiments

The human monocyte-like THP-1 cell line was obtained from ATCC (TIB-202) and cultivated in RPMI-media (ThermoFisher 72400047) with 10% FCS (ThermoFisher A5670502). Prior to stimulation, cells were seeded in 24-well plates, 3 x 10^5^ cells/well and differentiated into macrophage-like cells by treatment with Phorbol-12-myristate-13-acetate (Sigma, 524400, 180 nM) for three days. Following differentiation, cells were infected with *S. epidermidis*, MOI 1 for 6 h, with and without treatment with various concentrations of nicotinamide riboside (Sigma, SMB00907). Total RNA was isolated using the RNeasy kit according to manufacturer’s instructions (Qiagen, 7404). cDNA was synthesized from 280 ng of RNA using a High-Capacity cDNA Reverse Transcription Kit (Thermo Fisher, Waltham, MA, USA). Real-time quantitative PCR (RT–qPCR) was performed using the LightCycler 480 SYBR Green I Master kit on a LightCycler 480 (both Roche, Basel, Switzerland). Samples were analyzed in duplicate using cytokine-specific primers. Gene expression was normalized to the expression of beta-actin using the 2^-△△CT^ method. Data was normalized to controls. Statistical comparisons between treatments were performed by One-Way ANOVA and Fisher’s LSD.

#### Cord blood stimulation

Cord blood was collected from pig umbilical cords in EDTA tubes and processed immediately after birth. Blood was diluted 1:1 in RPMI and stimulated with 10^6^/ml UV-inactivated S.epidermidis, and 0.5 mM NR. Samples were incubated at 37 °C/5% CO_2_ for 5 h. Following incubation, supernatants were collected and cytokines measured using pig-specific ELISAs (see above).

### Statistics

#### Statistical analysis of COPSAC data

Associations between NAD levels at birth and infection risk during the first 3 years of life were analyzed using quasi-Poisson regression models with infection episodes as count outcomes. Time-to-first infection analyses were performed using Cox proportional hazards models. Associations between NAM levels and immune cell counts or cytokine responses were analyzed using general linear regression models. All analyses were performed in the total study population and stratified by sex. Sex interactions were assessed by including NAD/NAM-by-sex interaction terms in the relevant models. Analyses of birth NAD were adjusted for predefined covariates; fish oil and vitamin D trials, sex, social circumstances, maternal smoking during pregnancy, gestational age, birthweight, mode of delivery, neonatal hospitalization, maternal antibiotic use during pregnancy, maternal asthma, birth season, maternal age, maternal BMI, breastfeeding duration, and older siblings. Analyses of immune outcomes at 6 and 18 months were adjusted for trial randomization, sex, social circumstances, child BMI, and infections within the preceding 14 days, based on directed acyclic graphs and previous COPSAC analyses. Statistical analyses were performed using Stata (v14.2, Statacorp, USA), with statistical significance defined as P < 0.05.

#### Statistical analysis of in vivo experiments

Statistical analysis, data handling and visualization of in vivo experiments was performed in R (v4.5) with the lme4, emmeans, ggplot2, dplyr, tidyr, and patchwork packages. If model assumptions were not met, data were transformed (log or Box-Cox). For experiments consisting of several litters, a linear mixed-effects model including the fixed factors Timepoint (3 h, 6 h, 12 h, Euthanasia), Treatment group (CON-SE, NR-SE), sex and birthweight and the random intercept litter, was fitted. Fixed-effect significance was evaluated by F-tests with Satterthwaite degrees of freedom (lmerTest). If model assumptions were not met, the outcome was transformed (log, inverse, or Box–Cox as appropriate) and the analysis repeated. For individual comparisons, at each timepoint a separate mixed model (Outcome ∼ Group + Sex + Birthweight + (1|Litter)) was fitted. Pair-wise group contrasts were obtained from estimated marginal means with Tukey adjustment for multiple testing. Clinical data from animal experiments are presented as truncated violin plots with a line at the median and depiction of individual replicates.

In vitro experiments were analyzed using GraphPad prism 10.5.0. Typically, data were normalized to uninfected controls and compared using a ratio-paired t-test.

## Results

### NAD metabolism associates with reduced infection burden in female infants

Using the COPSAC2010 birth cohort with 700 children, we first investigated blood levels of NAD in early life and its association with infection burden. NAD levels in dry blood spots (DBS) at birth were significantly higher in females, compared to males (P_Sex_ < 0.001, **Fig. 1A**). We then correlated NAD levels at birth to the number of infection episodes in the first 36 months of life using a quasi-poisson regression adjusted for relevant confounders, yielding infection rate ratios (IRR) and corresponding 95% confidence intervals. Overall, there was a significant interaction between NAD and sex, with a negative correlation between total number of infections and NAD levels in females, but not in males (**Fig. 1B)**. For individual infections, the association between high NAD levels and reduced infection was significant in females for cold (IRR: 0.96 (0.93 – 0.99), P_NAD_ = 0.025), fever (IRR: 0.92 (0.88 – 0.98), P_NAD_ = 0.007) and gastroenteritis (IRR: 0.90 (0.82 – 0.99), P_NAD_ = 0.037), with a tendency for otitis media (IRR: 0.91 (0.82 – 1.00), P_NAD_ = 0.062). In males, higher NAD levels were by tendency associated with lower burden of tonsillitis (IRR: 0.81 (0.64 – 1.03), P_NAD_ = 0.092) but increased burden of otitis media (IRR: 1.08 (0.99 – 1.19), P_NAD_ = 0.097) and fever (IRR: 1.03 (0.98 – 1.09), P_NAD_ = 0.091). Time until the first reported infection was analyzed by a cox proportional hazard model using the same covariate adjustment yielding hazard ratios (HR) with corresponding 95% confidence interval. There was however no significant interaction between NAD and sex for any infection type, but higher NAD levels were overall associated with lower risk of fever (P_NAD_ = 0.049), an effect that was only significant in females (HR: 0.86 (0.75 – 0.97), P_NAD_ = 0.018) (**Sup. Fig. 1A**). Collectively, these clinical data suggest that the higher overall NAD levels observed in female infants around birth may contribute to their lower overall risk of common childhood infections, as compared to males.

**Figure 1.**
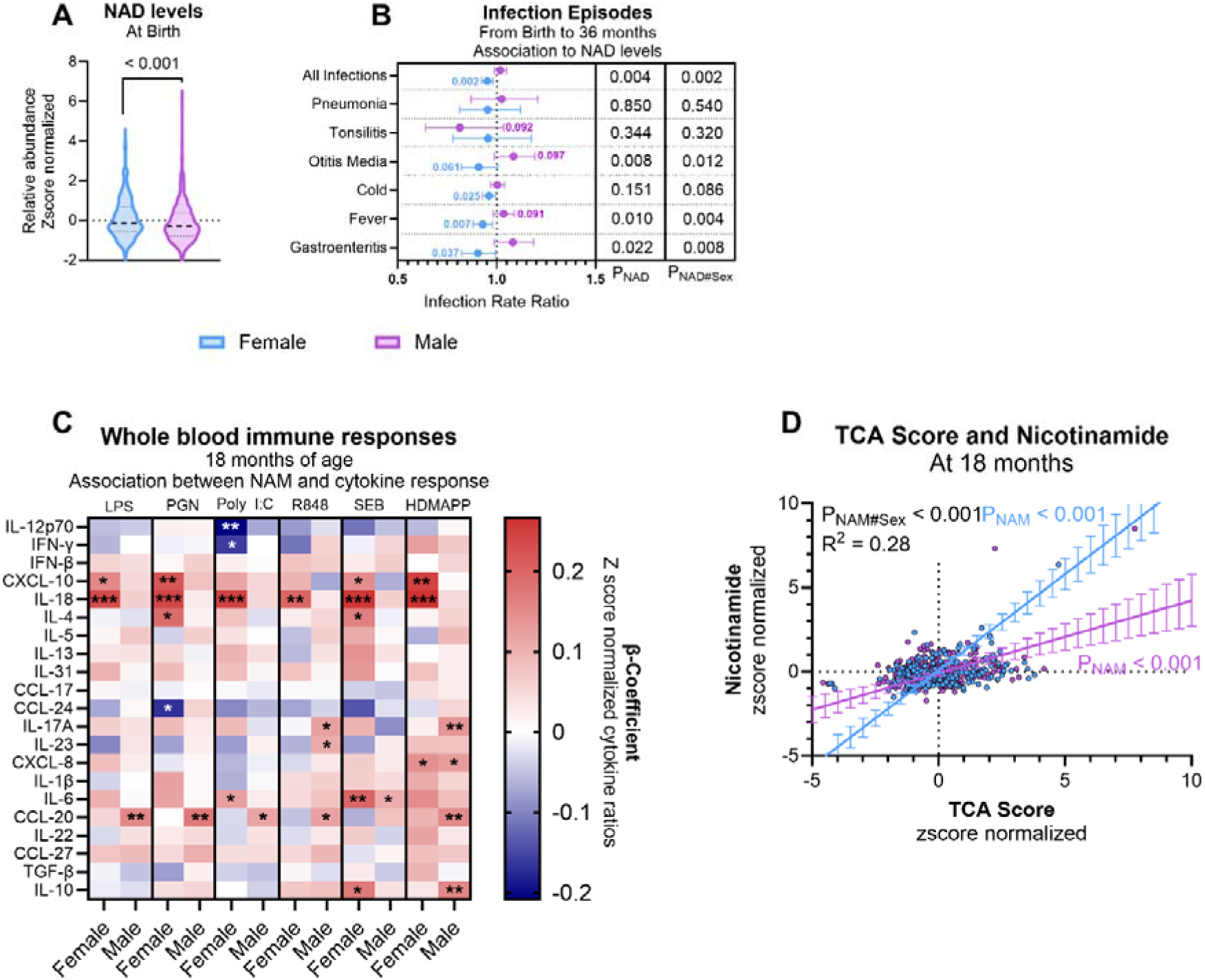
Sex-specific associations between NAD metabolism, immune responses, and childhood infections. **(A)** NAD levels in dry blood spots collected at birth, analyzed by non paramtetric test. **(B)** Associations between birth NAD levels and infection episodes from birth to 36 months, shown as sex-stratified infection rate ratios with 95% confidence intervals. P values indicate the overall NAD association and the interaction with sex. **(C)** Sex-stratified associations between nicotinamide (NAM) levels and ex vivo whole-blood cytokine responses to microbial and T-cell stimuli at 18 months. Colors indicate β-coefficients for z score-normalized cytokine ratios. *: P < 0.05, **: P < 0.01, ***: P< 0.001. **(D)** Association between TCA cycle score and nicotinamide levels at 18 months, stratified by sex. Blue indicates females and purple indicates males.

### NAD associates with improved immune function in females

To explore the association between NAD metabolism and immune function we used plasma and whole blood samples collected from children 6 and 18 months of age. In these samples only NAM, not NAD, could be detected. At 6 months of age, higher plasma NAM levels were associated with higher plasma levels of Interleukin- (IL) 1β and IL-8 (both P_NAM_ < 0.001), with no interaction with sex (**Suppl Fig. 1B**). At 18 months of age whole blood samples were stimulated with various microbial and T-cell stimuli for 24 h *ex vivo* and a range of cytokines measured (**Fig. 1D**). In females, higher NAM levels were positively associated with IL-18 responses across all stimulation conditions (all P_NAM_ < 0.01). This association was absent in males. Additional positive associations in females, but not males, were observed for CXCL10 after LPS, PGN, SEB, and HDMAPP stimulation; IL-4 after PGN and SEB stimulation; IL-6 after PolyI:C stimulation; and IL-10 after SEB stimulation. Higher NAM levels were also negatively associated with PolyI:C-induced IL-12p70 and IFN-γ responses, and with PGN-induced CCL24 in females only. In males, higher NAM was positively associated with CCL20 responses after all stimulations except SEB. Positive associations were also observed for IL-17A after R848 and HDMAPP stimulation, IL-23 after R848 stimulation, CXCL8 after HDMAPP stimulation, IL-6 after SEB stimulation, and IL-10 after HDMAPP stimulation. We also correlated the levels of NAM to fractions of different immune cell subsets measured by flow cytometry. Here, only levels of BDCA3+ dendritic cells were negatively associated with NAM in males (**Suppl Fig. 1C)**. All these results indicate that the plasma level of NAM is linked to distinct immune response patterns in females and males at 18 months of age.

We have previously shown that increased activity of the TCA cycle was strongly associated with a lower risk of common childhood infections using the same cohort.^19^ As such, we have previously calculated a child TCA score based on plasma levels of citrate, α-ketoglutarate, succinate, fumarate and malate. When analyzing the correlation between TCA score and NAM levels, these differed significantly by sex (P_NAM#Sex_ < 0.001), with females showing a more positive association between TCA score and NAM than males (**Fig. 1D**).

### Validation of findings in COPSAC2000

To independently verify our findings, we used a different birth cohort (COPSAC_2000_, n=386) with similar DBS based metabolomics performed at birth along with prospective recording of early life infections. In this independent cohort, levels of NAD were also higher in females compared to males (**Suppl Fig 1D**). Likewise, there was a significant interaction between NAD levels and sex for the overall risk of infections until 36 months of age (**Suppl Fig 1E**, P_NAD#Sex_ = 0.009). This was most evident for upper respiratory infections where NAD correlated with a lower risk in females (IRR: 0.93 (0.87 – 0.98), P_NAD_ = 0.005) but higher risk in males (IRR: 1.08 (1.03 – 1.13), P_NAD_ = 0.003). This confirms that the sex differences in early life NAD metabolism and infection risk are consistent in an independent cohort of children.

Collectively, our human data suggest that early-life NAD metabolism is linked to infection risk and immune function in a sex-dependent manner. Based on these findings, we hypothesize that female newborns may have greater mitochondrial metabolic capacity, which during infectious episodes could support a more controlled inflammatory response and reduce collateral tissue damage. Since NAD levels can be rapidly and safely increased by supplementation of the precursor NR, this may be a precision medicine approach to reducing the infection burden and improving outcomes in young females.

### NR treatment promotes bacterial clearance and survival in infected piglets in a sex-dependent manner

To explore the role of NAD metabolism during early life infection we used a well-established infection model in newborn piglets. Piglets were born by caesarean section at 90% gestation to mimic infection-prone newborns, fitted with umbilical arterial catheters, nourished by parenteral nutrition (PN) and infected with live *Staphylococcus epidermidis* as described in detail elsewhere.^31^ Prior to infection, piglets were stratified by sex and birthweight and allocated to receive either 200 mg/kg NR as intraarterial bolus (n = 18; 11 female, 7 male) or saline (n = 14; 8 female, 6 male).

Additionally, 200 mg/kg/18 h were administered via the PN solution over the course of infection (**Fig. 2A**). NR treatment significantly improved survival in female piglets, compared to female controls (4/11 to 7/8 survival, P = 0.046), an effect absent in males (**Fig. 2B**). Levels of hepatic NAD revealed that the NR administration was effective at boosting NAD production, with males reaching higher levels of NAD, compared to females (2200 nM to 1200 nM, P = 0.03, **Fig. 2C**). Bacterial burden in blood was also significantly affected by sex (P_Sex_ = 0.04), with a significant interaction with NR treatment (P_NR#Sex_ = 0.03). At euthanasia, NR-treated females showed lower bacterial burdens, compared to female controls (P = 0.02) and NR-treated males (P = 0.03). In contrast, blood bacterial burden in NR-treated males over all timepoints was significantly or by tendency increased, compared to male controls (**Fig. 2D**). For clinically relevant parameters there was a near significant interaction between NR treatment and sex for blood pH values across the experiment (P_NR#Sex_ = 0.06) with blood pH being lowest in NR treated males (**Fig. 2E)**. There was likewise a significant interaction between NR treatment and sex for blood carbon dioxide pressure (pCO_2_, P_NR#Sex_ < 0.01), with NR treatment reducing pCO_2_ in females, but increasing it in males (**Fig. 2F**). Accordingly, NR treatment by tendency did decrease oxygen saturation at later timepoints in males, but increased oxygen saturation in females (**Fig. 2G**). For lactate, there was a significant effect of sex, with males showing higher levels, compared to females (P_Sex_ = 0.03, **Fig. 2H**).

**Figure 2.**
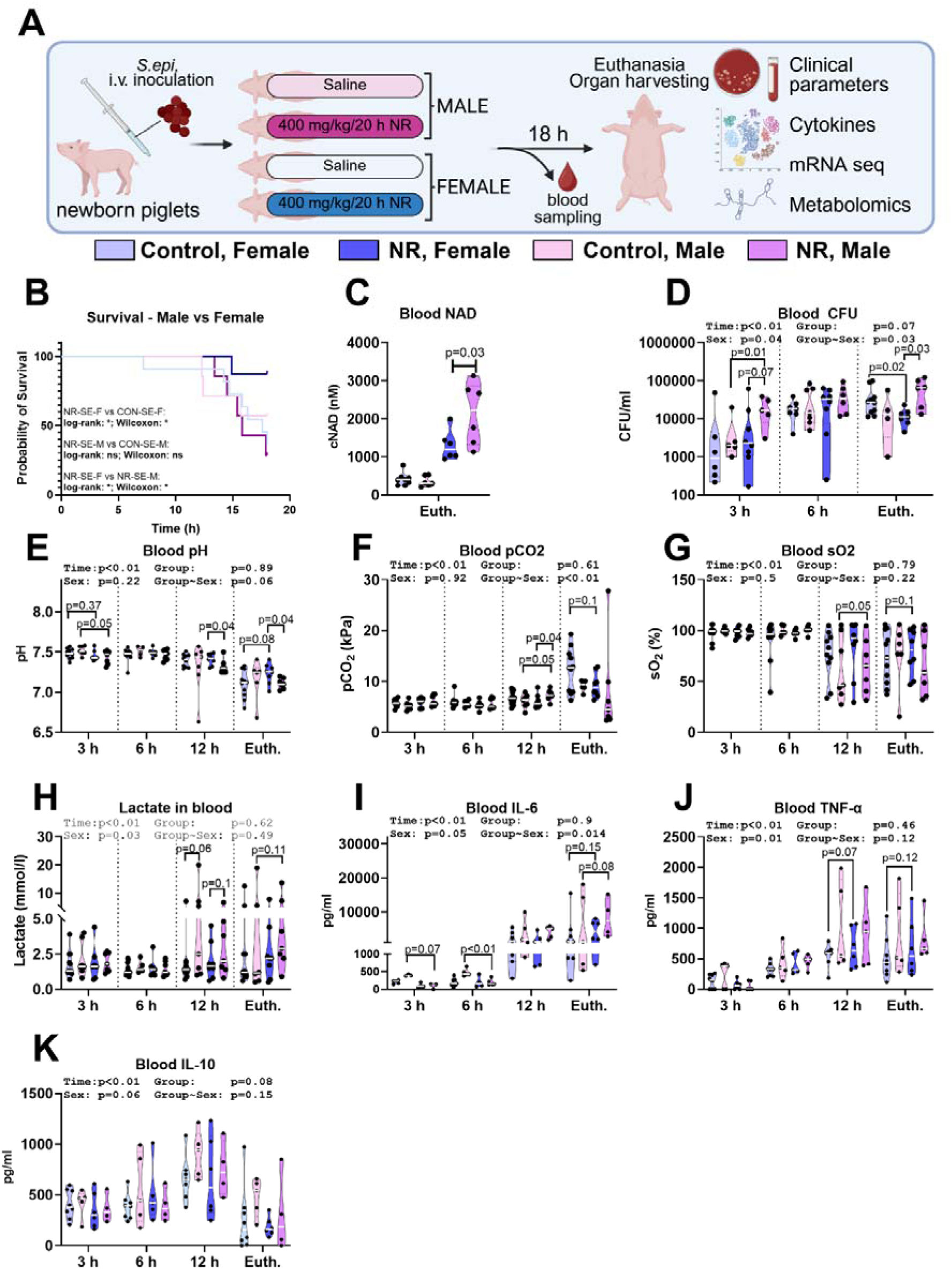
Clinical effects of NR treatment during neonatal infection. (A) Experimental setup. Male and female preterm neonatal piglets were infected with *S. epidermidis* by arterial infusion (10 CFU/kg). Piglets received either NR, administered as a 200 mg/kg bolus followed by 200 mg/kg over 18 h during infection (XX Female XX Male), or placebo(XX Female XX Male). Piglets were monitored for 18 h or until humane endpoint. (B) Survival, defined as time to humane endpoint, stratified by sex and NR treatement, analyzed by log-rank test. (C) Hepatic NAD levels at euthanasia analyzed by generalized linear model. (D) Blood bacterial burden (E–H) Blood gas parameters. (I–K) Plasma levels of IL-6, IL-10, and TNF-α., all analyzed by linear mixed-effects models.

To investigate how NR treatment affects immune responses during infection, we analysed plasma cytokine levels. There was a significant interaction between NR treatment and sex for IL-6 (P_NR#Sex_ = 0.01). In males, NR treatment reduced early IL-6 production, but increased levels during late infection, an effect not observed in females. (**Fig. 2I**). No main effect or significant interaction with sex was observed for TNF-a and IL-10 levels although by tendency, NR treatment increased levels of TNF-α in females 12 h after infection (**Fig.2 J, K**).

In summary, NR treatment during serious early life infection in pigs revealed marked sex-dependent effects. In females, NR treatment improved survival, reduced bacterial burdens and lowered lactate levels compared to NR-treated males, even though their hepatic NAD levels were lower.

### NR treatment drives disease tolerance in females

We have previously shown that improved survival during early-life infection is not determined solely by bacterial clearance or the magnitude of the inflammatory response but depends on the overall host defense strategy.^18^ Host defense during infection can be understood as a balance between disease resistance, the ability to eliminate pathogens from the system, and disease tolerance, the ability to maintain health despite increasing pathogen burden.^32,33^ We already observed that NR treated females showed lower bacterial burdens in blood, indicating an improvement in disease resistance in the later stages of the infection. To assess disease tolerance, we plotted blood pH (as a surrogate marker of overall health) against bacterial burden (**Fig. 3A**). A steeper negative slope indicates reduced tolerance, as pH drops more sharply with increasing bacterial load in the control group, female piglets depicted a steep negative slope at euthanasia (R = -0.44, P = 0.19), indicating a breakdown of disease tolerance in the later stages of the infection. Male controls maintained a shallow slope during the experiment, indicating maintenance of disease tolerance. In contrast, upon NR treatment, disease tolerance at euthanasia was observed in females (R_female_ = -0.11, P_female_ = 0.81), but not males (R_male_ = -0.82, P_male_ = 0.04). As such, the improved survival in NR treated females was accompanied by improvements in both disease resistance and tolerance allowing them to remove pathogens from the system while still maintaining normal organ function.

**Figure 3:**
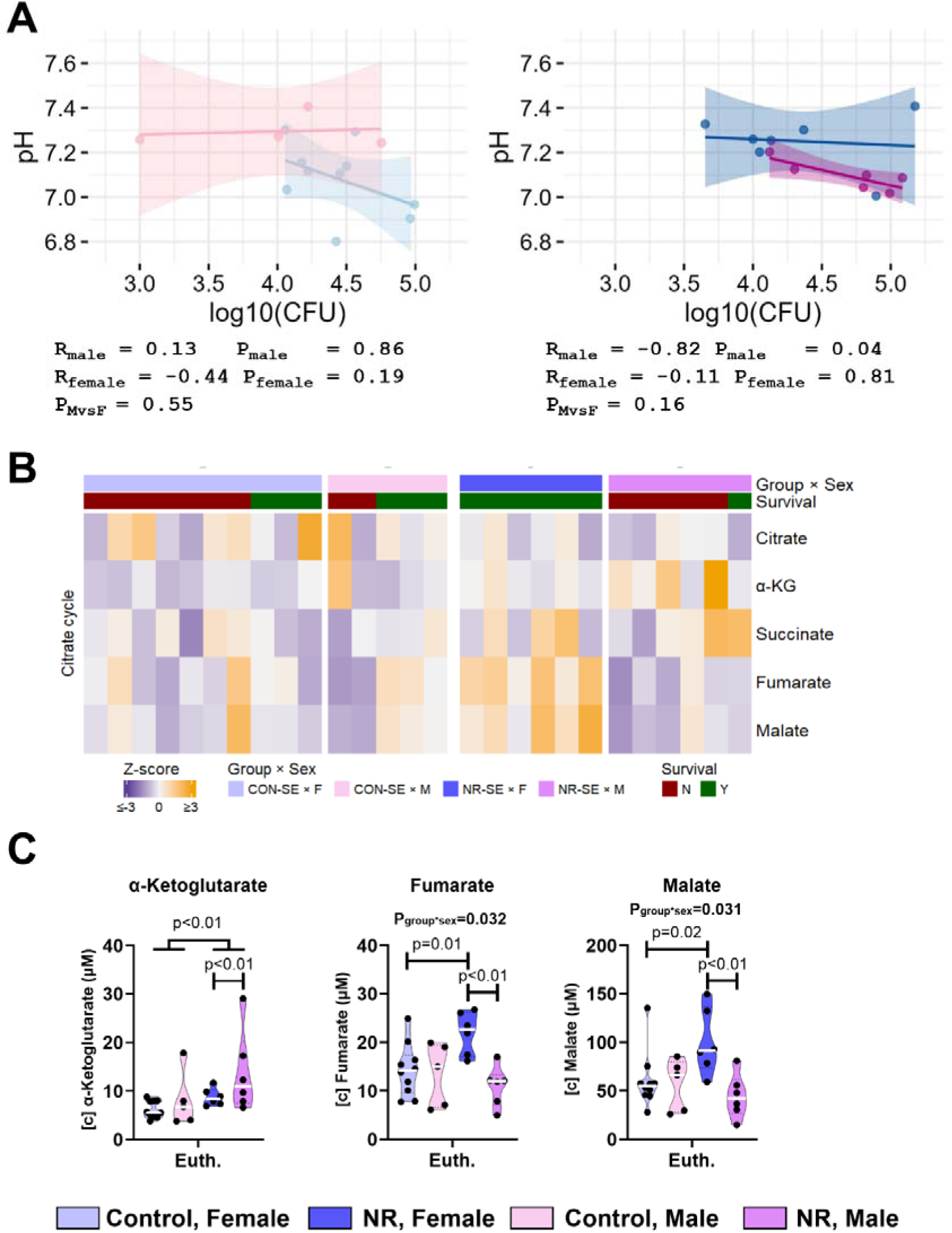
Targeted metabolomics and defense strategy of NR-treated piglets. A) Resistance-tolerance-analysis. At euthanasia, samples were collected from either male (pink/purple) or female (blue), infected piglets, treated with Nicotinamide riboside or placebo, as described above. A reaction-norm-analysis was performed to plot blood pH against the log_10_(CFU), with slopes calculated by linear regression. Slopes were compared using a sum-of-squares F-test. **B, C) Targeted LC-MS metabolomics of hepatic tissue** of infected, NR-treated piglets or infected controls was performed, focusing on metabolites associated with the TCA cycle. **B)** Z-score heatmap of metabolites associated with the TCA cycle. **C)** Intracellular concentrations of individual metabolites associated with the TCA cycle. (N=6-10).

### NR-treatment boosts mitochondrial function in a sex-dependent manner

We have also previously shown that improved infection survival is associated with modulated hepatic mitochondrial metabolism.^17–19,34^ To confirm that NR treatment was effective at improving mitochondrial function during infection, hepatic tissue sampled at euthanasia was subjected to targeted metabolomics, focusing on metabolites associated with the TCA cycle (**Fig. 3B-C**). NR treatment increased levels of α-ketoglutarate in both sexes, an effect more pronounced in males, compared to females. Further, NR treatment induced a significant increase in hepatic levels of the late TCA metabolites fumarate and malate in females (fumarate: 22 vs. 15 µM, P < 0.01; Malate: 80 vs. 55 µM, P = 0.02) (**Fig. 3C**), an effect that was absent in males. As such, NR treatment improved activity of the hepatic TCA cycle only in infected female piglets.

### NR treatment activates inflammatory pathways in females

Next, we assessed the hepatic transcriptomic response of female and male piglets to NR treatment during infection (**Fig. 4**). Kyoto Encyclopedia of Genes and Genomes Gene Set Enrichment Analysis (KEGG-GSEA) comparing female vs male NR-treated infected piglets revealed an activation of proinflammatory pathways in females (bacterial invasion of epithelial cells, false discovery rate (FDR) adjusted P = 0.01; mTOR signaling pathway, FDR adj. P = 0.07), compared to males (**Fig. 4A**). This was accompanied by an activation of hormonal and developmental signaling pathways (GnRH signaling pathways, FDR adj. P < 0.01; ErbB signaling pathway, FDR adj. P = 0.04). In contrast, comparing female vs male piglets in the control group showed a suppression of proinflammatory pathways (NF-kappa B signaling pathway, TNF signaling pathway) in females (**Fig. 4B**). Similarly, hormonal and developmental pathways were also suppressed in control females (growth hormone synthesis, secretion and action, FDR adj. P = 0.03; ErbB signaling pathway, FDR adj. P = 0.01).

**Figure 4:**
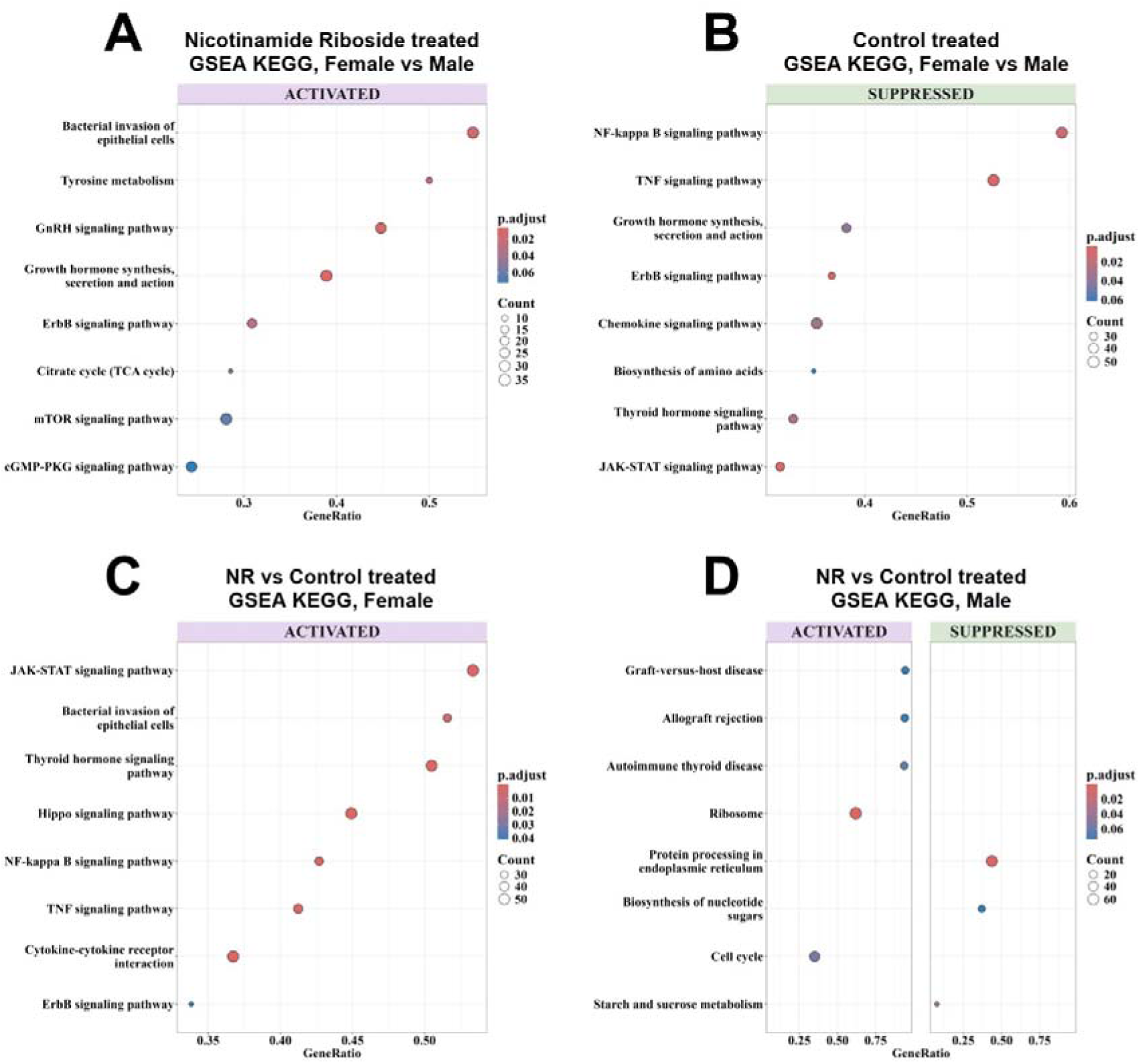
RNAseq analysis of NR-treated piglets. A-D) Gene-set enrichment analysis of differentially regulated genes in hepatic tissue, according to the KEGG database. **A)** Pathways identified in comparison of NR-treated, infected piglets, female vs male. Only activated pathways were identified **B)** Pathways identified in comparison of control-treated, infected piglets, female vs male. Only suppressed pathways were identified. **C)** Pathways identified in comparison of female, infected piglets, NR-treated vs control. Only activated pathways are depicted. **D)** Pathways identified in comparison of female, infected piglets, NR-treated vs control.

Comparing NR treated piglets vs controls within the female groups confirmed these findings: NR treatment resulted in an activation of inflammatory and hormonal pathways (JAK-STAT-signaling pathway, P < 0.01; thyroid hormone signaling pathway, P < 0.01), compared to controls (**Fig. 4C**). In contrast, within the males only, NR treatment had limited effects (**Fig. 4D**). Expression of individual genes of selected pathways was visualized by heatmaps. For “bacterial invasion of epithelial cells”, within the control groups, there are minor differences between males and females. Interestingly, the male non-survivors show a reduced gene expression, visualized by Z-scores. Upon NR-treatment, an activation of gene expression is observable in the females while gene expression in males is inhibited. A similar pattern is observable for “GnRH signaling pathway” (**Fig. S3**).

### Female-specific effects of NR treatment are conserved at a lower dose

Since hepatic NAD levels were higher in NR-treated female piglets than in males, we hypothesized that the NR dose used in our first animal study may have exceeded the beneficial range in males. We therefore tested whether a lower NR dose could improve infection outcomes in males while preserving the beneficial effects observed in females. In a second piglet infection experiment, using a design mimicking the previous study, piglets received an initial NR bolus of 50 mg/kg followed by 50 mg/kg over 20 h via parenteral nutrition (**Fig. 5**). Consistent with our previous findings, NR treatment significantly improved survival in infected females in this study, which had high overall mortality (CON: 0/11 survivors; NR: 4/11 survivors; P = 0.02). In contrast, survival was not significantly improved in males (CON: 2/12 survivors; NR: 4/13 survivors; P = 0.17) (**Fig. 5A**). NR treatment significantly affected bacterial burden in blood (P_NR_ = 0.01) (**Fig. 5B**). Although the NR-by-sex interaction was not significant (P_NR#Sex_ = 0.18), NR treatment reduced blood bacterial burden in females at 3 h (P = 0.02) and showed a near-significant trend towards a reduction at 6 h (P = 0.06) with no corresponding effect was observed in males. Effects on blood gas parameters were limited, with no significant changes in blood pH, lactate, pCO, or sO (**Fig. 5C–F**). Measurement of hepatic NAD+ and NADH+ revealed a moderate NAD-boosting effect of NR treatment, without significant differences between males and females. Overall, reducing the NR dose partially preserved its protective effects in infected female piglets but did not improve outcomes in males.

**Figure 5.**
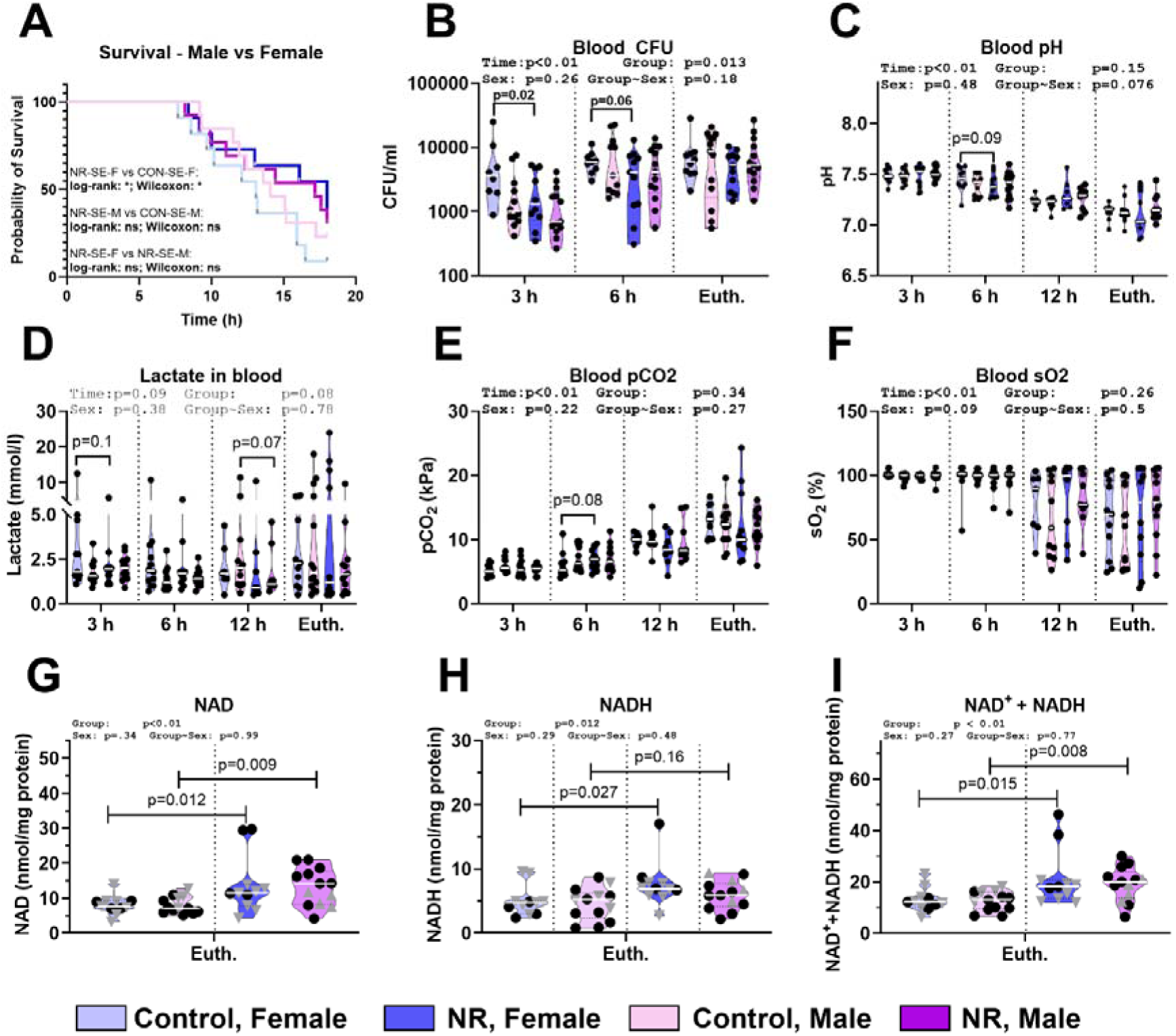
Clinical effects of lower-dose NR treatment during neonatal infection. Male and female preterm neonatal piglets were infected with S. epidermidis by arterial infusion (10 CFU/kg). Piglets received either NR, administered as a 50 mg/kg bolus followed by 50 mg/kg over 18 h during infection (11 Female 13 Male), or placebo (11 Female 13 Male). Piglets were monitored for 18 h or until humane endpoint. **(A)** Survival, defined as time to humane endpoint, stratified by sex and NR treatment, analyzed by log-rank test. **(B)** Blood bacterial burden. **(C-F)** Blood gas parameters. **(G-H)** Intracellular concentrations of NAD^+^, NADH and the sum of NAD+ and NADH, measured in hepatic tissue collected at euthanasia. All analyzed by linear mixed-effects models.

### NR treatment improves female immune function *in vitro*

Finally, we aimed to confirm a direct link between NR treatment and immune function *in vitro*. First, in male-derived PMA-differentiated THP-1 macrophages, we found a strong induction of cytokine gene expression following *S.epidermidis* infection, without any modulatory effect of NR treatment (**Fig. 6A-D**). In parallel, in cord blood from male and female newborn piglets, *in vitro* infection with UV-inactivated *S. epidermidis* induced secretion of TNF-α, but not IL-10, in the supernatant (**Fig. 6E-F**). When stratifying the response by sex, NR further potentiated infection-induced TNF-α secretion in female, but not in male blood.

**Figure 6:**
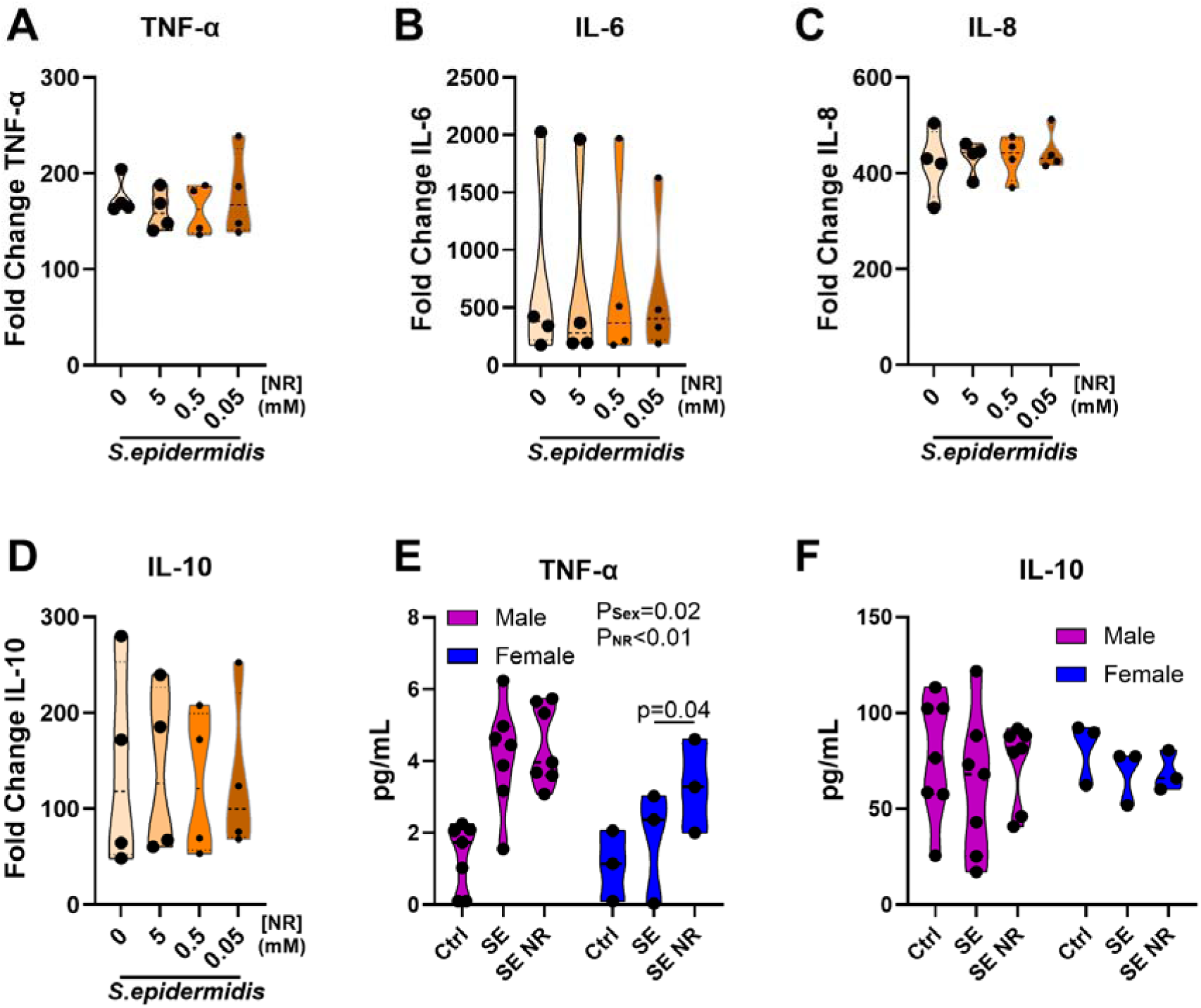
*In vitro* and *ex vivo* effects of NR treatment during infection. **A-D)** THP-1 monocyte like cells were differentiated using PMA for 3 days. Afterwards, cells were infected with S.epidermidis, MOI1 for 6 h, 37 C / 5% CO_2_, with and without treatment with indicated amounts of nicotinamide riboside (NR). Host RNA was isolated and gene expression determined by qPCR. (N=4) **E, F)** umbilical cord blood was collected from preterm-born piglets in EDTA tubes. Immediately after collection, blood was diluted 1:1 with RPMI and treated with either 10^6^ UV-inactivated S.epidermidis/ml (SE) and 0.5 mM NR solution or left untreated. Samples were incubated for 5 h 37 C / 5% CO_2_, supernatants were collected and cytokine secretion analysed by ELISA. (N=3-7; statistics: two-way ANOVA; t-test for individual comparisons).

## Discussion

Using two unique independent pediatric birth cohorts, we found that female infants had higher blood levels of NAD, compared to males. Further, we revealed that NAD levels at birth and NAM levels at 18 months of age were associated with reduced infection risk and improved circulating immune function only in females but not males. In certain conditions, NAD levels at birth were also associated with higher infection burden in males, although this effect was not consistent across the two cohorts. Then to demonstrate the causal effects of NAD metabolism, we employed a piglet model of early life infection and showed that NR supplementation during systemic infection improved bacterial clearance and infection survival outcomes only in females. These effects were mediated via enhanced hepatic mitochondrial metabolism, resulting in a balanced defense strategy improving both disease resistance and tolerance.

NR supplementation is a well-known intervention boosting NAD production^41,42^ and dysregulation of NAD metabolism has been observed during aging and various inflammatory and infectious diseases, including viral infection and sepsis^20^. Blood NAD levels decrease during aging, which may contribute to the increased infection risk commonly observed in the elderly^7,8,18,29^. However, clinical trials of NR supplementation have shown few beneficial outcomes.^35^

In humans, boosting NAD production using NR was demonstrated to improve COPD outcomes, indicated by decreased cytokine levels in sputum^45^. While some studies have shown that NR supplementation improves survival of septic rodents^36,37^, these studies did not stratify their results by sex and provided limited mechanistic insights. Likewise, very few NAD boosting interventions have been attempted in children. Supplementation of NAM (100mg/day) to infection prone infants in Tanzania reduced the risk of respiratory infections from 6 to 12 months of age, but effects were not powered and analyzed for sex.^38^

Now with the current study, we demonstrated across human cohorts, animal and *in vitro* models how NAD metabolism is connected to infection risk, infection responses and immune function in a sex-dependent manner. NAD levels in humans and NR interventions in piglets and immune cells *in vitro* were consistently connected to improved infection outcomes in females, without affecting or, to a degree, worsening outcomes in males. To our knowledge, this is the first report that an immunometabolic intervention benefits one sex while harming another in the context of early life infections. As we observed these effects over two different experimental doses of NR, this indicates that the results may be mediated by sex-diverging downstream effects of NAD, rather than purely by differences in NAD biosynthesis pathways. While some studies showed an anti-inflammatory effect of NR supplementation^35,39^, others reported increased inflammatory response connected to elevated blood NAD levels, e.g. during atherosclerosis^40^. In the piglet study, we demonstrated that NR supplementation induced a time-dependent modification of cytokine release, diminishing cytokine response in the early phase of infection but improving the response in the later phase. These kinetic effects may explain diverging findings in previous studies.

Importantly, we found that NAM levels in the blood of 18-month-old children correlated with TCA cycle activity, with a higher correlation coefficient in female vs. male children. In parallel, in infected piglets NR treatment was more efficient in enhancing the two key metabolites fumarate and malate at the end of TCA cycle in the liver. NAD levels can be significantly influenced by PARP1 and SIRT3 pathways, but the outcomes are highly divergent: NF-kB-induced inflammatory responses by PARP1 but enhanced TCA cycle activity mediated by SIRT3.^41,42^ Prioritization of these pathways may, directly or indirectly, be dictated by sex-associated genes, thereby mediating these sex-stratified infection outcomes. Further, the female-preferred effect of NAD on TCA cycle metabolites was observed even though NR treatment increased hepatic levels of NAD stronger in males than in females. This indicates either a more efficient uptake of NR, a more efficient conversion of NR into nicotinic acid adenine dinucleotide (NAAD) or less consumption of the produced NAD in males.

Beside improved disease resistance (indicated by less bacterial density or better bacterial clearance), NR-treated, infected female piglets also utilized a disease tolerance defense strategy (analyzed by reaction-norm analysis) across the whole infection period, a process that is typically associated with improved disease outcomes^18,32,43^. Hepatic transcriptomics of these NR-treated animals also revealed a higher expression of inflammatory and hormonal signaling pathways in females. These results were inverted in the infected control group, where females depicted a lower expression of inflammatory pathways. Importantly, this does not come at the cost of impaired antimicrobial function, as NR-treated females exhibited a reduced bacterial burden in blood, compared to control-treated females or NR-treated males. These effects appear to be correlated with enhanced inflammatory responses and immune responses to *ex vivo* challenges in COPSAC human infants. The sex-specific improvement of infection outcomes persisted in a second experiment where a lower dose of NR was used for treatment. Here, NR treatment increased intra-hepatic levels of NAD+ and NADH to similar levels in males and females. Still, NR treatment improved survival in females only, confirming our previous observation and indicating that the sex-divergent effect of NR treatment is not driven by differences in NR utilization. *In vitro* and *ex vivo* cell stimulation assays further confirm the sex dependency of NR on immune function. In the male-derived^44^ THP-1 cell line, NR treatment during infection had no effect on cytokine expression. In contrast, treatment of umbilical cord blood resulted in a sex-dependent effect on cytokine release, with NR inducing secretion of TNF-α only in females. These data further support a model of direct immune-cell associated factors as mediators of the sex-dependent effects we observed.

There are limitations to our study. The infection episodes recorded in the COPSAC cohort were mostly mild to medium severity and recorded differently in the two cohorts. In contrast, our animal experiments modeled an acute, severe infection. Still, our *in vivo* result confirmed improved immune and antimicrobial functions upon NR treatment, thereby providing empirical evidence for the observational findings from the COPSAC cohort. From a translational aspect, the dose of NR supplementation in newborn animals led to blood NAD levels beyond physiological concentrations often aimed for in existing clinical trials.^45,46^ Regardless, this work provides proof-of-concept of sex-dependent effects of NAD metabolism in infections that can pave the way for clinical trials testing sex-stratified NAD-boosting interventions in infection-sensitive infants and children.

## Acknowledgements

We gratefully acknowledge Thomas Thymann, Kristina Larsen, Malene Skovsted Cilieborg, Britta Karlsson, and Jane Connie Povlsen for their valuable support with the animal experiments and laboratory analyses. The experimental work was supported by the Novo Nordisk Foundation to DNN (Grant No. NNF22OC0078747). OB received support from the BRIDGE–Translational Excellence Programme at the Faculty of Health and Medical Sciences, University of Copenhagen, funded by the Novo Nordisk Foundation (Grant Nos. NNF23SA0087869 and NNF20SA00643). BC and TW were supported by the European Research Council through the European Union’s Horizon 2020 research and innovation program (Grant No. 946228). Mass spectrometry-based metabolomics was conducted at the Metabolomics Platform, Novo Nordisk Center for Basic Metabolic Research, University of Copenhagen. We thank Raissa Novais Rodrigues and Muyao Xi from the Metabolomics Platform for their technical assistance and expertise.

## Supplementary Material

**Supplementary Figure 1.**
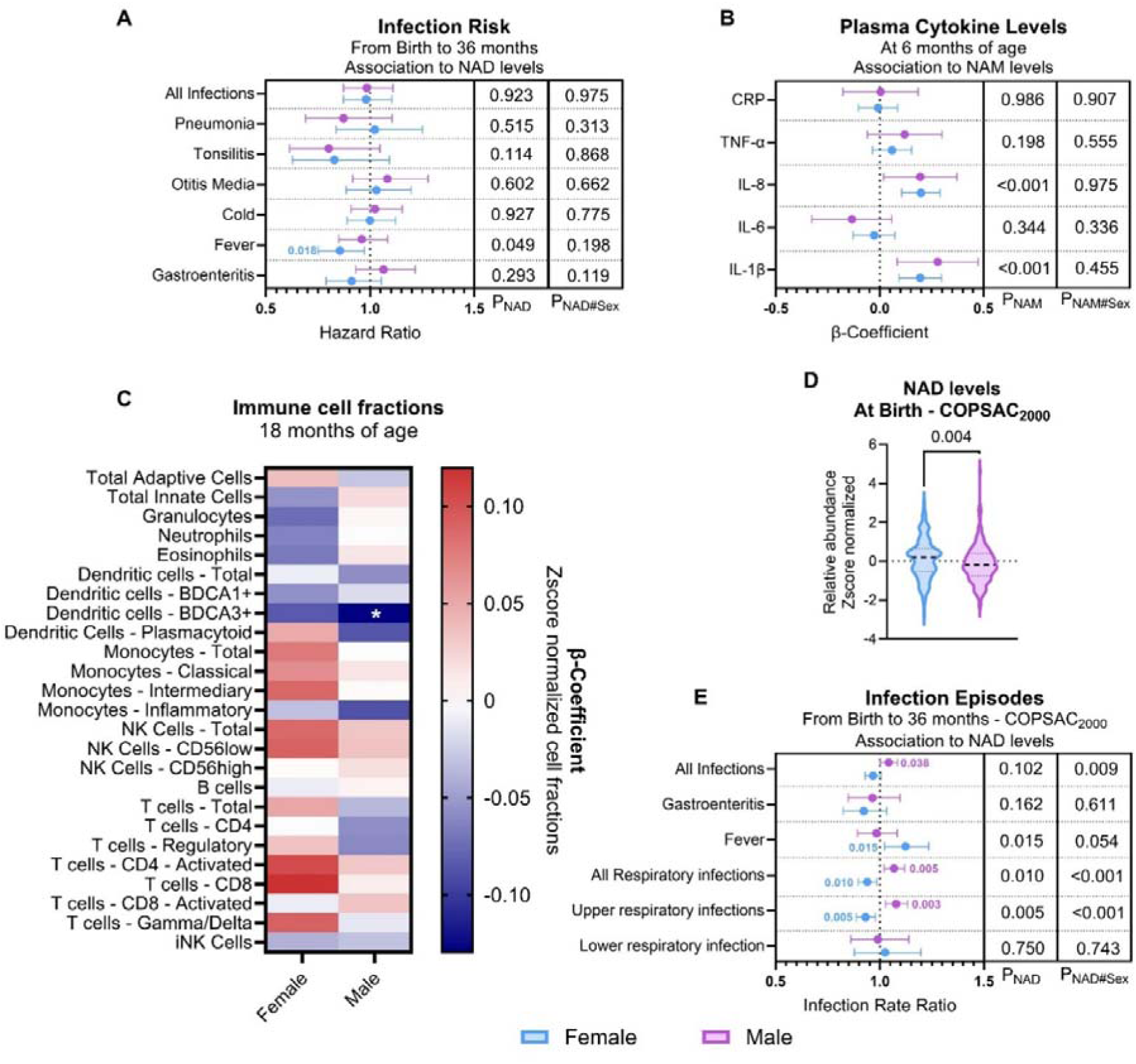
Additional sex-specific associations between NAD metabolism, immune phenotypes, and childhood infections. (A) Associations between NAD levels at birth and time to first infection from birth to 36 months, shown as sex-stratified hazard ratios with 95% confidence intervals and analyzed using Cox proportional hazards models. (B) Associations between nicotinamide (NAM) levels and plasma cytokine levels at 6 months of age, shown as sex-stratified β-coefficients with 95% confidence intervals. (C) Sex-stratified associations between NAM levels and circulating immune cell fractions at 18 months. Colors indicate β-coefficients for z score-normalized cell fractions. *: P < 0.05. (D) NAD levels at birth in the independent COPSAC2000 cohort, stratified by sex. (E) Associations between birth NAD levels and infection episodes from birth to 36 months in COPSAC2000, shown as sex-stratified infection rate ratios with 95% confidence intervals. P values indicate the overall NAD or NAM association and the NAD/NAM-by-sex interaction. Blue indicates females and purple indicates males.

**Supplemental Figure 2:**
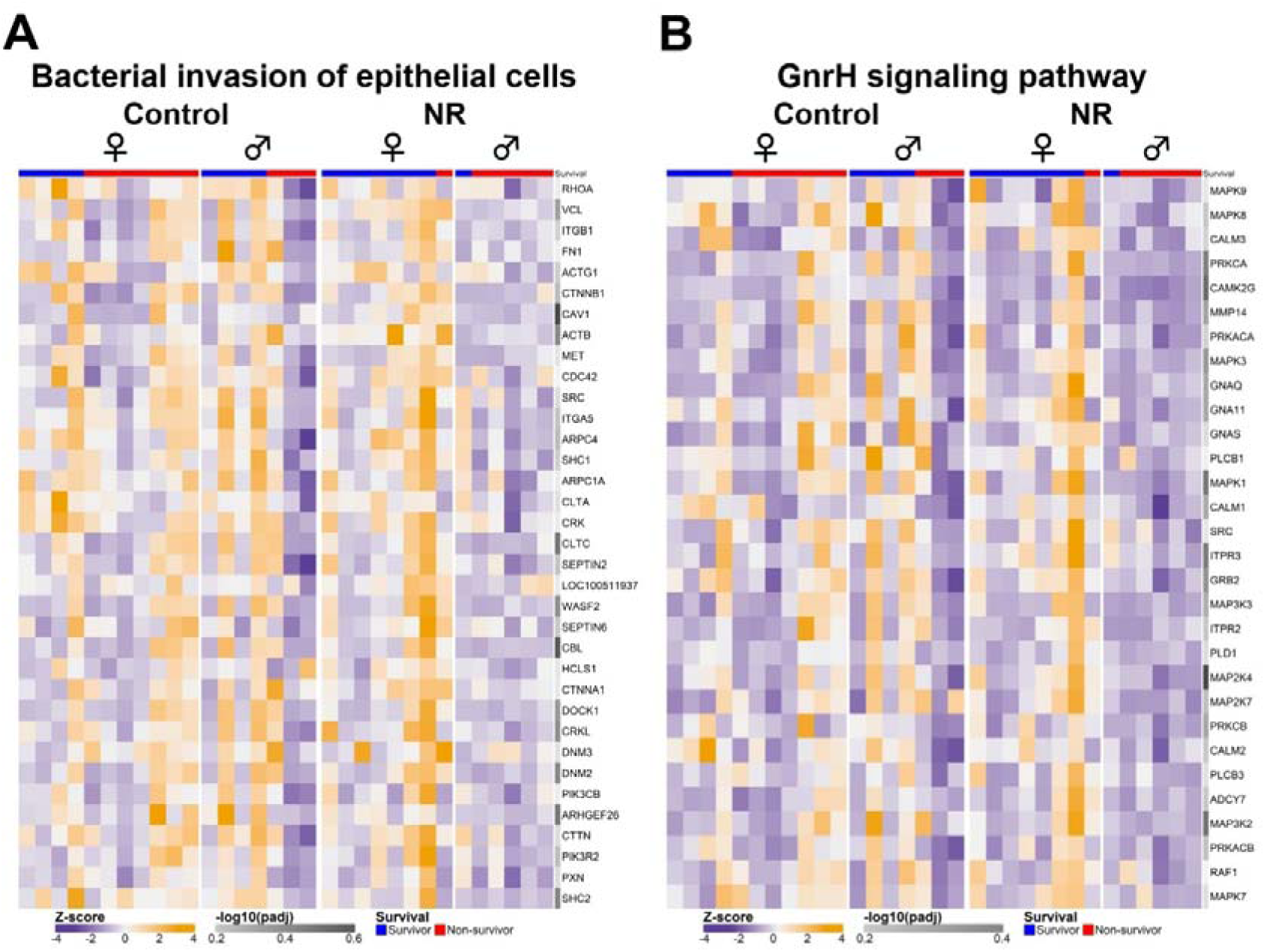
Extended transcriptomics following NR supplementation. **A,B)** Heatmap of genes associated with the KEGG pathways “Bacterial invasion of epithelial cells” **(A)** and “GnRH signaling pathway” **(B)**. Expression is depicted as Z-scores, calculated based on vst-transformed gene expression in all samples. N = 7-10.

## Supplemental Tables

**Supplemental Table 1:** Retention time of target metabolites.

| Precursor.Name | Label.Type | Precursor.Formula | Precursor.Adduct | Precursor.m/z | Retention.Time |
| --- | --- | --- | --- | --- | --- |
| Citric acid |  | C<sub>24</sub>H<sub>23</sub>N<sub>9</sub>O<sub>10</sub> | [M-H]- | 596.1495 | 6.89 |
| Malic acid |  | C<sub>16</sub>H<sub>16</sub>N<sub>6</sub>O<sub>7</sub> | [M-H]- | 403.1008 | 5.54 |
| Succinic acid |  | C<sub>16</sub>H<sub>16</sub>N<sub>6</sub>O<sub>6</sub> | [M-H]- | 387.1059 | 5.83 |
| Fumaric acid |  | C<sub>16</sub>H<sub>14</sub>N<sub>6</sub>O<sub>6</sub> | [M-H]- | 385.0902 | 6.40 |
| Pyruvic acid |  | C<sub>15</sub>H<sub>14</sub>N<sub>6</sub>O<sub>5</sub> | [M-H]- | 357.0953 | 7.83 |
| Pyruvic acid |  | C<sub>15</sub>H<sub>14</sub>N<sub>6</sub>O<sub>5</sub> | [M-H]- | 357.0953 | 7.35 |
| D-glucose |  | C<sub>12</sub>H<sub>17</sub>N<sub>3</sub>O<sub>7</sub> | [M-H]- | 314.0994 | 3.54 |
| Alpha-ketoglutarate |  | C<sub>23</sub>H<sub>21</sub>N<sub>9</sub>O<sub>8</sub> | [M-H]- | 550.1440 | 8.42 |
| Alpha-ketoglutarate | heavy | C<sub>23</sub>H<sub>19</sub>H'<sub>3</sub>N<sub>9</sub>O<sub>8</sub> | [M-H]- | 554.1692 | 8.40 |
| Citric acid | heavy | C<sub>21</sub>C'<sub>3</sub>H<sub>23</sub>N<sub>9</sub>O<sub>10</sub> | [M-H]- | 599.1596 | 6.89 |
| Malic acid | heavy | C<sub>12</sub>C'<sub>4</sub>H<sub>16</sub>N<sub>6</sub>O<sub>7</sub> | [M-H]- | 407.1142 | 5.53 |
| Succinic acid | heavy | C<sub>16</sub>H<sub>12</sub>H'<sub>4</sub>N<sub>6</sub>O<sub>6</sub> | [M-H]- | 391.1310 | 5.80 |
| Fumaric acid | heavy | C<sub>16</sub>H<sub>10</sub>H'<sub>4</sub>N<sub>6</sub>O<sub>6</sub> | [M-H]- | 389.1153 | 6.44 |

## References

1. Kyvsgaard, J. N. et al. Key risk factors of asthma-like symptoms are mediated through infection burden in early childhood. Journal of Allergy and Clinical Immunology 153, 684–694 (2024).

2. Grüber, C. et al. History of respiratory infections in the first 12 yr among children from a birth cohort. Pediatric Allergy and Immunology 19, 505–512 (2008).

3. Brustad, N. et al. Burden of Infections in Early Life and Risk of Infections and Systemic Antibiotics Use in Childhood. JAMA Netw. Open 8, e2453284 (2025).

4. Hibbert, J. E., Currie, A. & Strunk, T. Sepsis-induced immunosuppression in neonates. Front. Pediatr. 6, (2018).

5. Kariniotaki, C. et al. Neonatal Sepsis: A Comprehensive Review. Antibiotics 14, 6 (2024).

6. Preau, S. et al. Energetic dysfunction in sepsis: a narrative review. Ann. Intensive Care 11, 1– 21 (2021).

7. Stearns-Kurosawa, D. J., Osuchowski, M. F., Valentine, C., Kurosawa, S. & Remick, D. G. The pathogenesis of sepsis. Annu. Rev. Pathol. 6, 19–48 (2011).

8. Wasyluk, W. & Zwolak, A. Metabolic Alterations in Sepsis. J. Clin. Med. 10, 2412 (2021).

9. Murthy, S., Godinho, M. A., Guddattu, V., Lewis, L. E. S. & Sreekumaran Nair, N. Risk factors of neonatal sepsis in India: A systematic review and meta-analysis. PLoS One 14, (2019).

10. Bech, C. M. et al. Risk factors for neonatal sepsis in Sub-Saharan Africa: a systematic review with meta-analysis. BMJ Open 12, e054491 (2022).

11. Muenchhoff, M. & Goulder, P. J. R. Sex Differences in Pediatric Infectious Diseases. J. Infect. Dis. 209, S120 (2014).

12. Klein, S. L. & Flanagan, K. L. Sex differences in immune responses. Nature Reviews Immunology 2016 16:10 16, 626–638 (2016).

13. Forsyth, K. S., Jiwrajka, N., Lovell, C. D., Toothacre, N. E. & Anguera, M. C. The conneXion between sex and immune responses. Nat. Rev. Immunol. 24, 487–502 (2024).

14. Klabunde, B. et al. NAD+ metabolism is a key modulator of bacterial respiratory epithelial infections. Nature Communications 2023 14:1 14, 1–16 (2023).

15. Groth, B., Venkatakrishnan, P. & Lin, S. J. NAD+ Metabolism, Metabolic Stress, and Infection. Front. Mol. Biosci. 8, 686412 (2021).

16. Katsyuba, E., Romani, M., Hofer, D. & Auwerx, J. NAD+ homeostasis in health and disease. Nat. Metab. 2, 9–31 (2020).

17. Bæk, O. et al. Altered hepatic metabolism mediates sepsis preventive effects of reduced glucose supply in infected preterm newborns. Elife 13, (2025).

18. Wu, Z. et al. Regulation of host metabolism and defense strategies to survive neonatal infection. Biochimica et Biophysica Acta (BBA) - Molecular Basis of Disease 1870, 167482 (2024).

19. Wu, Z. et al. Harnessing the glycolysis-TCA cycle axis to boost host defense against neonatal infection. EMBO Mol. Med. 10.1038/S44321-026-00463-Z (2026) doi:10.1038/S44321-026-00463-Z.

20. Zheng, M., Schultz, M. B. & Sinclair, D. A. NAD+ in COVID-19 and viral infections. Trends Immunol. 43, 283–295 (2022).

21. Rago, D. et al. Characteristics and Mechanisms of a Sphingolipid-associated Childhood Asthma Endotype. Am. J. Respir. Crit. Care Med. 203, 853–863 (2021).

22. Rago, D. et al. Fish-oil supplementation in pregnancy, child metabolomics and asthma risk. EBioMedicine 46, 399 (2019).

23. Sumner, L. W. et al. Proposed minimum reporting standards for chemical analysis Chemical Analysis Working Group (CAWG) Metabolomics Standards Initiative (MSI). Metabolomics 3, 211 (2007).

24. Thysen, A. H. et al. Distinct immune phenotypes in infants developing asthma during childhood. Sci Transl Med 12, (2020).

25. Love, M. I., Huber, W. & Anders, S. Moderated estimation of fold change and dispersion for RNA-seq data with DESeq2. Genome Biol. 15, 1–21 (2014).

26. R: A Language and Environment for Statistical Computing. https://www.gnu.org/copyleft/gpl.html.

27. Bioconductor - EnhancedVolcano. https://bioconductor.org/packages/release/bioc/html/EnhancedVolcano.html.

28. Yu, G., Wang, L. G., Han, Y. & He, Q. Y. clusterProfiler: an R package for comparing biological themes among gene clusters. OMICS 16, 284–287 (2012).

29. Gu, Z., Eils, R. & Schlesner, M. Complex heatmaps reveal patterns and correlations in multidimensional genomic data. Bioinformatics 32, 2847–2849 (2016).

30. Jensen, J. B., et al. A randomized placebo-controlled trial of nicotinamide riboside and pterostilbene supplementation in experimental muscle injury in elderly individuals. JCI Insight 7, (2022).

31. Muk, T., Brunse, A., Henriksen, N. L., Aasmul-Olsen, K. & Nguyen, D. N. Glucose supply and glycolysis inhibition shape the clinical fate of Staphylococcus epidermidis–infected preterm newborns. JCI Insight 7, (2022).

32. Medzhitov, R., Schneider, D. S. & Soares, M. P. Disease tolerance as a defense strategy. Science 335, 936–941 (2012).

33. Weis, S. et al. Metabolic Adaptation Establishes Disease Tolerance to Sepsis. Cell 169, 1263–1275.e14 (2017).

34. Bæk, O. et al. Diet Modulates the High Sensitivity to Systemic Infection in Newborn Preterm Pigs. Front. Immunol. 11, (2020).

35. Damgaard, M. V. & Treebak, J. T. What is really known about the effects of nicotinamide riboside supplementation in humans. Sci. Adv. 9, eadi4862 (2023).

36. Hong, G. et al. Administration of nicotinamide riboside prevents oxidative stress and organ injury in sepsis. Free Radic. Biol. Med. 123, 125–137 (2018).

37. Zhao, G. J. et al. SUPPLEMENTATION WITH NICOTINAMIDE RIBOSIDE ATTENUATES T CELL EXHAUSTION AND IMPROVES SURVIVAL IN SEPSIS. Shock 60, 238 (2023).

38. DeBoer, M. D. et al. Effect of scheduled antimicrobial and nicotinamide treatment on linear growth in children in rural Tanzania: A factorial randomized, double-blind, placebo-controlled trial. PLoS Med. 18, e1003617 (2021).

39. Elhassan, Y. S. et al. Nicotinamide Riboside Augments the Aged Human Skeletal Muscle NAD+ Metabolome and Induces Transcriptomic and Anti-inflammatory Signatures. Cell Rep. 28, 1717 (2019).

40. Wang, Y. J. et al. NAD+ boosting increases atherosclerotic plaques and inflammation in Apoe knockout mice. Atherosclerosis 404, (2025).

41. Houtkooper, R. H., Pirinen, E. & Auwerx, J. Sirtuins as regulators of metabolism and healthspan. Nat. Rev. Mol. Cell Biol. 13, 225–238 (2012).

42. Bai, P. et al. PARP-1 inhibition increases mitochondrial metabolism through SIRT1 activation. Cell Metab. 13, 461 (2011).

43. Harbeson, D., Francis, F., Bao, W., Amenyogbe, N. A. & Kollmann, T. R. Energy Demands of Early Life Drive a Disease Tolerant Phenotype and Dictate Outcome in Neonatal Bacterial Sepsis. Front. Immunol. 9, 408524 (2018).

44. Tsuchiya, S. et al. Establishment and characterization of a human acute monocytic leukemia cell line (THP-1). Int. J. Cancer 26, 171–176 (1980).

45. Norheim, K. L. et al. Effect of nicotinamide riboside on airway inflammation in COPD: a randomized, placebo-controlled trial. *Nat*. Aging 4, 1772–1781 (2024).

46. Martens, C. R. et al. Chronic nicotinamide riboside supplementation is well-tolerated and elevates NAD+ in healthy middle-aged and older adults. Nature Communications 2018 9:1 9, 1–11 (2018).

